# Empirical evidence and robustness of clock models with spikes

**DOI:** 10.64898/2026.08.11.744144

**Authors:** Haobo Yuan, Ewan Ciuffi, Timothy G. Vaughan, Daniele Silvestro, Tanja Stadler

## Abstract

Time-calibrated phylogenies provide information on past macroevolutionary history. Time calibration can be obtained from fossil ages or node calibrations in combination with a clock model describing evolutionary rates. Relaxed clocks, which allow rates of evolution to vary across lineages, are widely used in phylogenetic research but lack a mechanistic link between rate variation and the evolutionary process. A recently developed class of clock models attempts to introduce biological mechanisms by coupling speciation events with spikes of evolutionary change. However, their empirical support and overall impact on phylogenetic inference remain underexplored. Here, we evaluate the support for spike clock models across a range of empirical datasets and use simulations to quantify the effects of model misspecification and missing data across clock models. We find that spike clock models are supported as the best-fitting model in six of the seven datasets analyzed, suggesting widespread evidence of a punctuated mode of evolution. Although the choice of clock models does not strongly affect the resulting divergence time estimates, spike clock models tend to give more constrained uncertainty intervals of speciation and extinction rate estimates in some empirical analyses. We interpret this as the consequence of information transfer from sequence evolution into the inferred branching process. Our simulations show that a general clock model that incorporates both branch-specific clock rates and spikes is the most robust across simulated datasets. In particular, models with spikes are robust to missing data, capable of accurately estimating speciation and extinction rates even when fossil data is completely absent. In summary, we highlight here that evolutionary spikes leave a detectable signal in the alignment data, and correctly accounting for them leads to improved estimates of the branching parameters and tree topologies.

## 2 Introduction

Estimation of time-calibrated phylogenies is key to understanding evolution in absolute timescales. As genetic and morphological distances among taxa are a product of time and evolutionary rates, clock models have been developed to allow us to disentangle their contributions and reconstruct phylogenetic trees with branch lengths representing divergence times. Relaxed clock models assign a unique substitution rate to each branch and are commonly used to accommodate rate heterogeneity across lineages because of their widespread empirical support against constant-rate strict clock models. Early relaxed clock models were formulated in a mechanistic way, assuming that substitution rates are inherited from ancestral to descendant lineages (Thorne *et al*. 1998; Kishino *et al*. 2001; Rannala and Yang 2007). These so-called autocorrelated clock models imply that closely related species share similar substitution rates, as a result of the transmission of heritable characteristics that determine rates of molecular evolution (*e.g.*, generation time and longevity) through descent (Sanderson 2002; Sanderson 2003; Bromham 2020). Later, Drummond *et al*. (2006) proposed uncorrelated relaxed clocks, which assume that rates of evolution are independently drawn from a common distribution across branches. Under these models, substitution rates on adjacent lineages are not expected to be more similar than those on distantly related lineages. The motivation behind uncorrelated relaxed clocks is that the rate autocorrelation assumption may break down in studies of widely divergent species and low taxon sampling, where patterns of rate inheritance will diminish over long evolutionary timescales, or closely related lineages, where the rate variation across branches will be dominated by stochasticity rather than inheritance (Drummond *et al*. 2006). Despite their wide popularity in Bayesian dating analyses due to statistical and computational convenience, uncorrelated relaxed clocks as well as other recent clock model developments have been criticized for being phenomenological, failing to explicitly account for macroevolutionary processes that may cause rate heterogeneity (Lartillot *et al*. 2016; Ho 2020; Bromham 2020).

More recently, another class of clock models has been developed to explicitly account for spikes of evolutionary change that occur at speciation events (*i.e.*, punctuated evolution; Eldredge and Gould 1972), thus attempting to reincorporate mechanistic interpretations for evolutionary rate variation (Manceau *et al*. 2020; Janzen *et al*. 2022; Douglas *et al*. 2025). Under spike clock models, the evolutionary change is determined by evolutionary spikes occurring at observed and unobserved speciation events. These events are, in turn, estimated based on the birth-death process. As a result, spike clock models are the first clock models to couple the likelihood of sequence alignments with the diversification process in phylogenetic reconstruction, due to this association of spikes with speciation events. Importantly, spike clock models allow for simultaneous reconstruction of time-calibrated phylogenies and detection of punctuated change, whereas previous efforts have largely focused on inferring punctuated evolution from phenotypic traits of fossil time-series data (Hunt 2007; Hunt *et al*. 2008; Hunt 2008; Hunt 2012; Grey *et al*. 2008; Hopkins and Lidgard 2012; Voje 2016; Voje *et al*. 2020; see reviews by Duran-Nebreda *et al*. 2024; Hunt *et al*. 2025) or based on fixed topologies, such as regression analysis (Webster *et al*. 2003; Pagel *et al*. 2006; Rabosky 2012; Rabosky and Adams 2012; Rabosky *et al*. 2013; Surya *et al*. 2023) and phylogenetic comparative methods (Bokma 2002; Bokma 2008; Mattila and Bokma 2008; Bokma 2010; Landis *et al*. 2013; Bartoszek 2014; Duchen *et al*. 2017; Mitov *et al*. 2020; Bastide and Didier 2023).

The empirical support and actual impact of spike clock models on phylogenetic inference in macroevolutionary studies remain to be fully characterized. In this study, we explore the extent to which spike clock models are supported compared to relaxed clock alternatives across a range of empirical datasets that include different proportions of extant and extinct taxa and combinations of molecular and morphological data. Following this, we perform extensive simulations to establish the robustness and potential biases of using spike or standard relaxed clock models in face of model misspecification and missing data. As spike clock models develop into a new generation of more mechanistic ways to reconstruct evolution and phylogenetic histories, our results characterize the potential benefits and limitations of this framework.

## 3 Methods

### 3.1 Clock models

We evaluated four clock models (Figure 1a), namely 1) the strict clock (hereafter: STR), where the clock rate is the same across the whole tree, 2) the uncorrelated log-normal relaxed clock (RLX; Drummond *et al*. 2006), where each branch is assigned its own clock rate from a log-normal distribution, 3) the uncorrelated log-normal clock with gamma spikes (RLX+S; Douglas *et al*. 2025), in which the sequence mutates instantaneously at each speciation event with rates drawn from a gamma distribution, and 4) the previously untested strict clock with spikes (STR+S), where rate variation only occurs at speciation events in the form of spikes. We used the STR+S model to assess to what extent spikes can absorb rate variation generally captured by relaxed clocks and its relative fit when applied to empirical data. In this study, we will refer to the latter two models that allow for spikes as “spike clock models” collectively. Further details on spike clock models are provided in the Supplementary Information.

**Figure 1:**
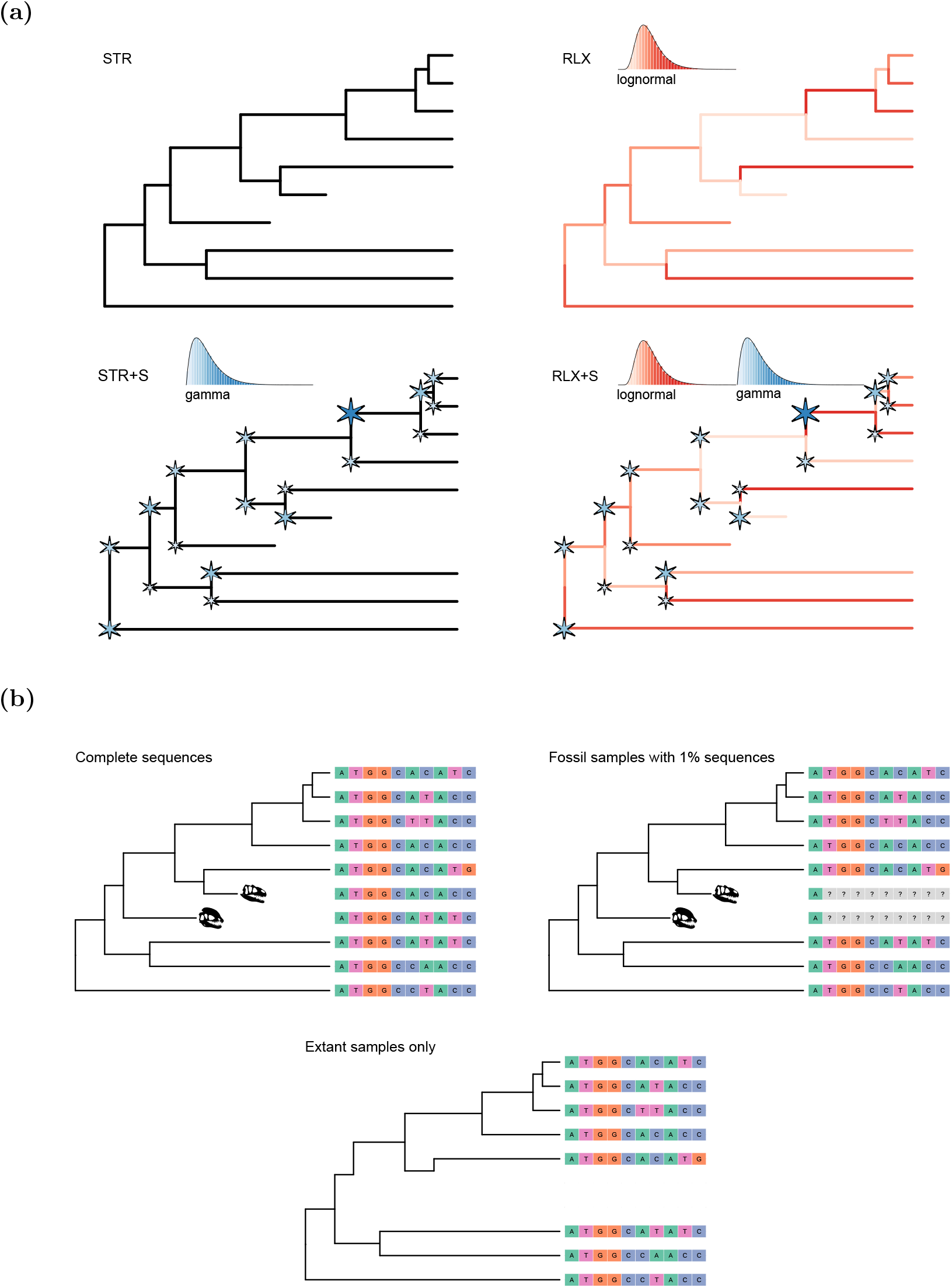
Schematic diagram showing (a) four clock models applied to empirical datasets and (b) three inference scenarios investigated in the simulation study with decreasing data completeness.

We compared the performance of clock models across a range of empirical and simulated datasets. Specifically, we first evaluated their relative fit on empirical datasets and their impact on estimated parameters. Then, we analyzed simulated alignments, focusing on the three clock models that allow for rate variation (*i.e.*, RLX, STR+S, and RLX+S) to investigate their robustness to model misspecification and missing data.

### 3.2 Analysis of empirical data

We analyzed seven published empirical datasets, namely Cephalopoda (Whalen and Landman 2022), Canidae (Stolz *et al*. 2025), Platyrrhini (Silvestro *et al*. 2019), Proboscidea (Baleka *et al*. 2022), Spheniscidae (penguins; Gavryushkina *et al*. 2017), Soricdae (shrews; Yuan *et al*. 2024), and Caviomorpha (Carrillo *et al*. 2026). These datasets differ in their number of taxon samples, size and type of comparative data, and evolutionary time scales (Table S1). All datasets have been originally analyzed using the RLX model. Here, we reanalyzed them under the four clock models mentioned above, while keeping the original inference model set-up and prior choices where possible. Minor modifications were made to better reflect current prior knowledge and improve Markov chain Monte Carlo (MCMC) mixing. All settings are detailed in the Supplementary Information. The cephalopod dataset was previously analyzed by Whalen and Landman (2022) using a spike model (RLX+S), but without incorporating information about the sampling fraction of extant species included in the analysis, thus implicitly assuming full sampling (*ρ* = 1). However, estimates of living cephalopod diversity indicate a much lower value of *ρ* = 0.03375 (Lindgren *et al*. 2004). To demonstrate the potential impact of accounting for incomplete sampling, we repeated the analyses under both assumptions (*ρ* = 1 or 0.03375).

Spike clock models involve two additional parameters: the mean spike size (*S_µ_*) and the shape parameter (*S_α_*) that describes the gamma distribution of relative spike sizes. We used log-normal priors for both: *S_µ_* ∼ LogNormal(mean = 0.01*, σ* = 1.2) and *S_α_* ∼ LogNormal(mean = 2*, σ* = 0.5). We ran five replicates of each analysis and combined the final log and tree files after discarding the first 10% of samples as burn-in. To compare parameter estimates between clock models, we focused on speciation and extinction rates and total branch length and calculated the median and 95% highest posterior density (HPD) interval of the posterior samples. We also compared the inferred tree topologies under different clock models by calculating quartet distances of the sampled topologies and projecting the calculated distances to a two-dimensional space using multi-dimensional scaling (Mongiardino Koch *et al*. 2026). We performed these analyses on a sample of 1000 posterior trees.

We performed model testing on the four clock models using stepping-stone sampling to infer marginal likelihoods (Xie *et al*. 2011). To quantify the uncertainty around marginal likelihood estimates, we ran ten replicates and summarized them as the mean and standard deviation. To compare relative model fit, we calculated the log Bayes factors (*i.e.*, differences in log marginal likelihood estimates). We followed the criteria defined by Kass and Raftery (1995) and used threshold of one, three, and five log units as positive, strong, and very strong support for the model with the higher marginal likelihood.

### 3.3 Simulation and inference under different clock models

We used simulations to assess the performance of different clock models in face of model misspecification and missing data. We focused on the estimated trees (topology and total branch lengths) and branching parameters (speciation and extinction rates). We simulated sequence alignments under three clock models that allow for rate variation (RLX, STR+S, and RLX+S) and performed inference using the same three clock models, resulting in nine combinations of simulation and inference models. This allowed us to investigate the performance of the three clock models under the true and misspecified inference conditions. We additionally applied three different treatments to the sequence alignments to mimic the effect of incomplete data (Figure 1b).

We used FossilSim (Barido-Sottani *et al*. 2019) to simulate trees with 30 extant and a variable number of fossil tips under the fossilized birth-death process (Heath *et al*. 2014) with parameters drawn from the following distributions: speciation rate *λ* ∼ *U*(0.1, 1), turnover rate 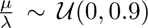, fossil sampling rate *ψ* ∼ U(0.1, 0.3), and extant sampling proportion *ρ* = 1. We discarded trees with sampled ancestors on the root edge and only retained trees with 5 − 150 fossil tips. The resulting 100 simulated trees had a median number of 57 tips (95% range: 36 − 129) and a median tree height of 11.03 units of time (95% range: 3.07 − 38.99). A summary of the simulated trees is shown in Table S10. We simulated branch-specific clock rates using an uncorrelated relaxed clock model, with mean clock rate *µ_c_* ∼ U(0.001, 0.01) and standard deviation of relative clock rates *σ_r_* ∼ Γ(*α* = 5*, θ* = 0.05). We simulated branch-specific spikes with mean spike size *S_µ_* ∼ LogNormal(mean = 0.05*, σ* = 0.8) and shape parameter of relative spike sizes *S_α_*∼ LogNormal(mean = 2*, σ* = 0.5). For each clock model, we simulated alignments of 2500 sites using the HKY model with four-category gamma-distributed across-site rate variation (Hasegawa *et al*. 1985; Yang 1994) with the transition/transversion ratio drawn from Exp(mean = 5) and gamma shape parameter from Exp(mean = 1).

We tested three scenarios which decreased progressively in data completeness: (1) extant and fossil samples were included with their complete sequences, (2) fossils only preserved the first 25 sites (1%) of their sequence data, or (3) fossil samples were removed from the dataset and only extant samples were used for inference (Figure 1b).

For inference, we used the fossilized birth-death model as implemented in the BDMM-Prime package (Vaughan and Stadler 2025) as the tree prior. To control for the effect of uncertainty in tree age estimation, we conditioned the tree prior on the root age and fixed the tree height to the true values. We further used broad and uninformative priors for the following parameters: speciation rate *λ* ∼ Exp(mean = 1), turnover rate 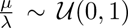, fossil sampling rate *ψ* ∼ Exp(mean = 1), mean clock rate *µ_c_* ∼ Exp(mean = 1), and mean spike size *S_µ_*∼ Exp(mean = 0.05). For all other parameters, we applied the prior distributions used in simulations.

We focused on speciation and extinction rates to assess the inferred branching dynamics as well as total tree length to assess the time calibration in the estimated trees. For these parameters, we calculated the relative error of posterior medians compared to the true values. We computed the relative error of speciation and extinction rates using the sum of the two rates as the denominator. To assess model performance in estimating tree topologies, we calculated the normalized Robinson-Foulds distances between the sampled posterior trees and the true topology for each analysis (Robinson and Foulds 1981). We discarded the first 10% of posterior samples as burn-in.

## 4 Results

### 4.1 Clock model comparison in empirical data

All seven empirical datasets showed evidence of across-branch rate variation, with the STR model consistently achieving lower support than all alternative models (Table S2; Figure 2). These results also indicate that a model with a homogeneous clock rate and heterogeneous spikes at speciation events (STR+S) fits empirical data overwhelmingly better than the STR model, in some cases even outperforming the relaxed clock alternatives (*e.g.*, canids, platyrrhines, and proboscideans; Figure 2c-e). Furthermore, spike clock models were supported in six of the seven clades analyzed.

**Figure 2:**
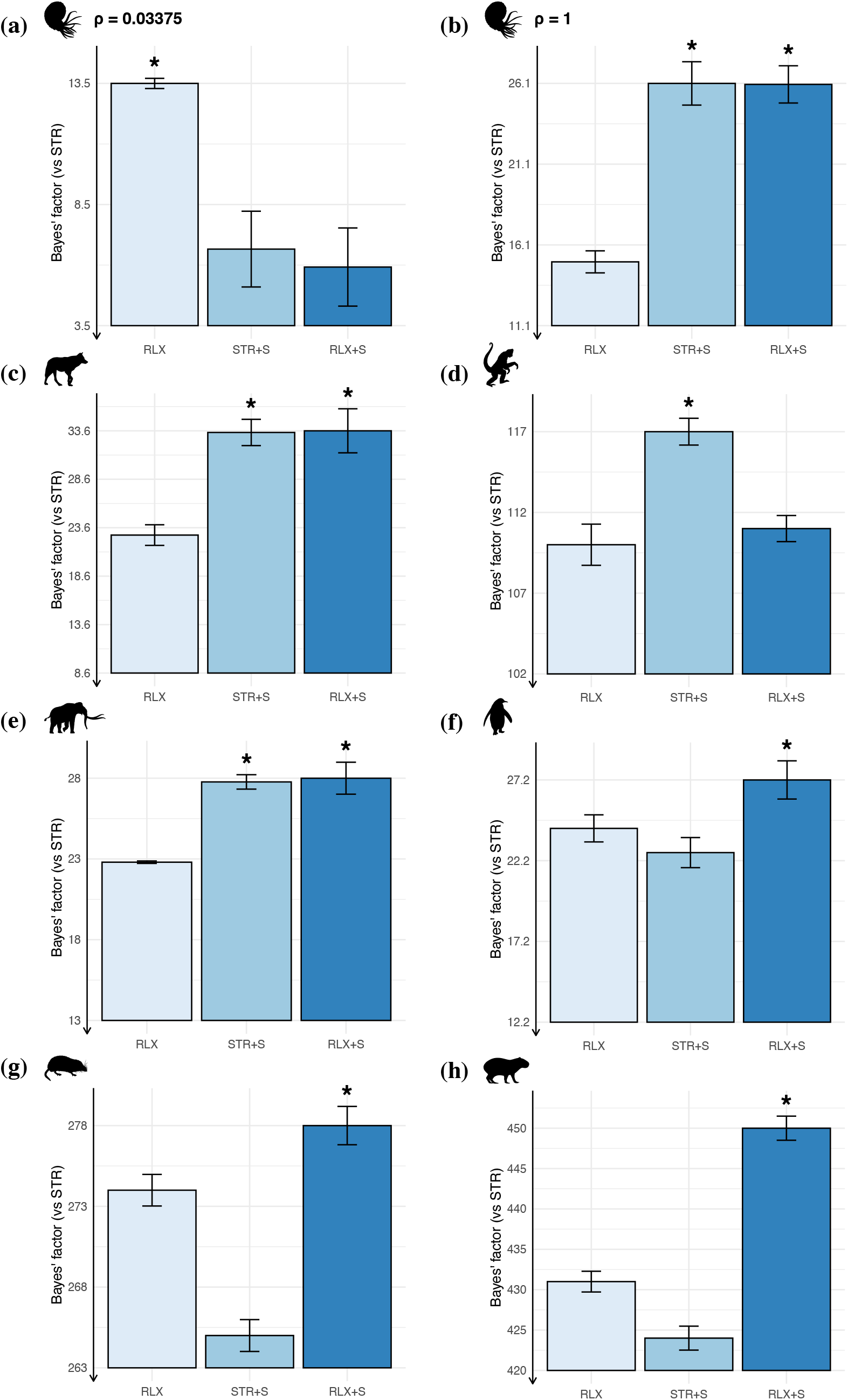
Mean Bayes factor estimates with standard deviation against the strict clock (STR) model under three clock models that allow for rate variation. (a) Cephalopods with empirical *ρ* sampling probability. (b) Cephalopods with complete *ρ* sampling probability. (c) Canids. (d) Platyrrhines. (e) Proboscideans. (f) Penguins. (g) Shrews. (h) Caviomorphs. The best-fitting model is indicated by an asterisk.

In the cephalopod analyses, we found that accounting for incomplete taxon sampling had a major impact on the selected clock model. Indeed, under the (erroneous) assumption of complete sampling for extant cephalopods (*ρ* = 1; Figure 2b), we find overwhelming support for spike clock models over RLX. However, when we account for incomplete sampling (*ρ* = 0.03375; Figure 2a), the RLX model is strongly favored over spike clock models. All other datasets showed strong evidence for one or both spike clock models over RLX. While the two spike clock models obtained similar support in canids and proboscideans (Figure 2c and 2e), the best fitting model for platyrrhines was the STR+S model (Figure 2d). In the remaining datasets, namely penguins, shrews, and caviomorphs, the most parameter-rich model RLX+S was unanimously favored over the alternatives (Figure 2f-h). Thus, all empirical datasets except for the cephalopod, which has extremely low taxon sampling, show support for an evolutionary process with spikes at branching events.

### 4.2 Branching parameter and tree estimates

Given the overwhelming support for models that allow for across-branch rate variation, we focused on comparing the estimates of branching parameters and trees between the RLX, STR+S, and RLX+S models. While the tree length estimates were generally consistent across models, we found substantial differences in speciation and extinction rate estimates between RLX and the two spike clock models in several groups, namely cephalopods, canids, proboscideans, shrews and, to a lower extent, caviomorphs (Figure 3a, b, d, f, g). In these cases, the spike clock models produced consistently lower speciation and extinction rate estimates with narrower 95% HPD intervals than those estimated by the RLX model. For instance, in cephalopods, where the best fitting model was RLX, the median speciation and extinction rate estimates under the spike clock models were 56% − 59% lower and the 95% HPD intervals were 60% narrower than those under RLX (Table S3; Figure 3a). A similar pattern is however observed also in datasets where spike clock models received strong support against RLX, such as canids and proboscideans. In canids, the median speciation and extinction rate estimates under the spike clock models were 20% − 21% lower and the 95% HPD intervals 53% − 54% narrower than those inferred under RLX (Table S4; Figure 3b). In proboscideans, the median speciation and extinction rate estimates were 55% − 57% lower and the 95% HPD intervals 82% − 83% narrower under the spike clock models (Table S6; Figure 3d). In shrews, the best-supported RLX+S model estimated median speciation and extinction rates that were 21% and 28% respectively lower and HPD intervals that were 36% than RLX, whereas the alternative spike clock model STR+S produced 37% and 47% lower median speciation and extinction rates and nearly 60% narrower HPD intervals (Table S8; Figure 3f). The difference was less pronounced in caviomorphs, where the median speciation and extinction rate estimates under the spike clock models were only 7% − 12% lower than under RLX (Table S9; Figure 3g). In contrast, the speciation and extinction rate estimates in platyrrhines and penguins were stable across all three models (Tables S5, S7; Figure 3c, e).

**Figure 3:**
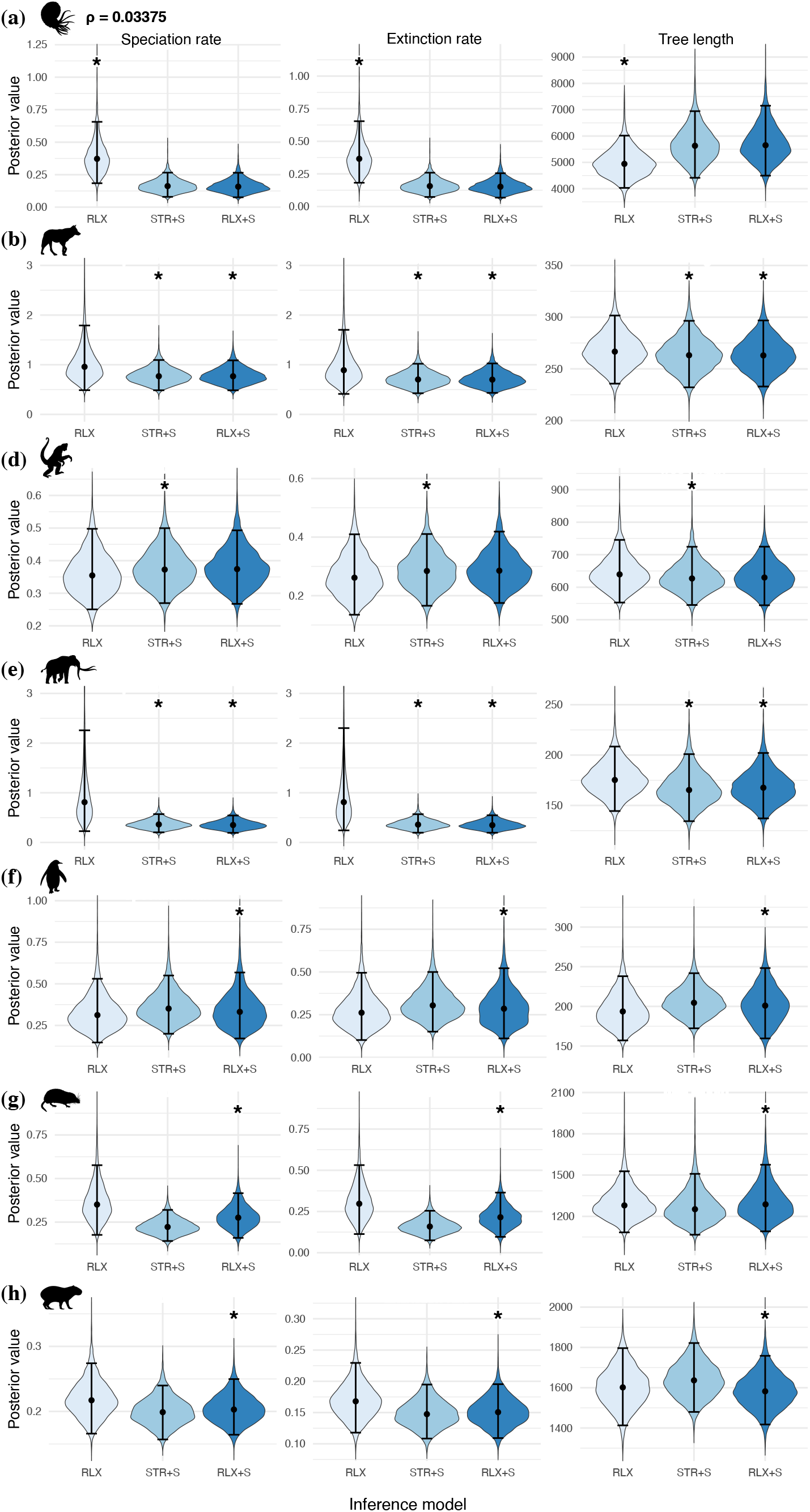
Posterior estimates of speciation rate, extinction rate, and tree length under three clock models. (a) Cephalopods with empirical *ρ* sampling probability. (b) Canids. (c) Platyrrhines. (d) Proboscideans. (e) Penguins. (f) Shrews. (g) Caviomorphs. The best-fitting model is indicated by an asterisk. The point represents the median and the error bar represents the 95% HPD interval.

We do not find evidence for substantial differences in estimated total branch lengths among the seven datasets, as the estimated medians and 95% HPD intervals are overall consistent among the three clock models (Tables S3-S9; Figure 3). While the spike clock models produced higher median tree length estimates (∼ 14%) in cephalopods (Table S3; Figure 3a), the relative differences in estimated medians and HPD widths between the spike clock models and RLX were negligible (0% − 6% and 0% − 14%, respectively) in the other clades (Tables S4-S9; Figure 3b-g).

Additionally, RLX and spike clock models produced tree topologies that overlapped substantially in the two-dimensional tree space of quartet distances, indicating that despite discrepancies in the estimated branching parameters, the three models inferred highly similar phylogenetic relationships. In contrast, topologies inferred by the STR model occupied a distinct tree space in some of the clades, especially in penguins, shrews, and caviomorphs (Figure S1e-g).

### 4.3 Simulation study

Analyses of simulated datasets showed that the branching parameters and tree length were generally most accurately estimated under the scenario where complete sequences were available for both extant and fossil samples (Table S11; Figure 4a). When the datasets were analyzed under the correct clock model, the inferred parameters and tree lengths were unbiased across simulations both with and without spikes, with median relative errors centred around 0. The RLX+S model, the most general clock in our analyses, produced unbiased estimates across datasets (median relative errors between −0.01 and 0.03), including those generated under the STR+S and RLX models. In contrast, the STR+S model generated overestimated speciation and extinction rates (∼ 6%) and slightly underestimated total tree length (∼ 2%) when the simulated data do not contain spikes. When data were simulated with spikes, the two spike clock models gave similarly accurate estimates of branching parameters and tree lengths, whereas the RLX model underestimated speciation rate (∼ 9%−10%) and extinction rate (∼ 3%−4%) and overestimated total tree length by 13% on average. The inferred tree topologies were highly accurate overall when we used complete sequence data for inference, with median normalized Robinson-Foulds distances ranging between 0.01 and 0.07 (Table S14; Figure 4a). When analyzing data simulated without spikes, all three clock models produced similarly accurate tree topologies (median distance: 0.07). Using the RLX model to analyze data simulated under spike clock models resulted in an increase in the topological error of the estimated trees (median distance: 0.04) compared to spike clock models (median distance: 0 − 0.01).

**Figure 4:**
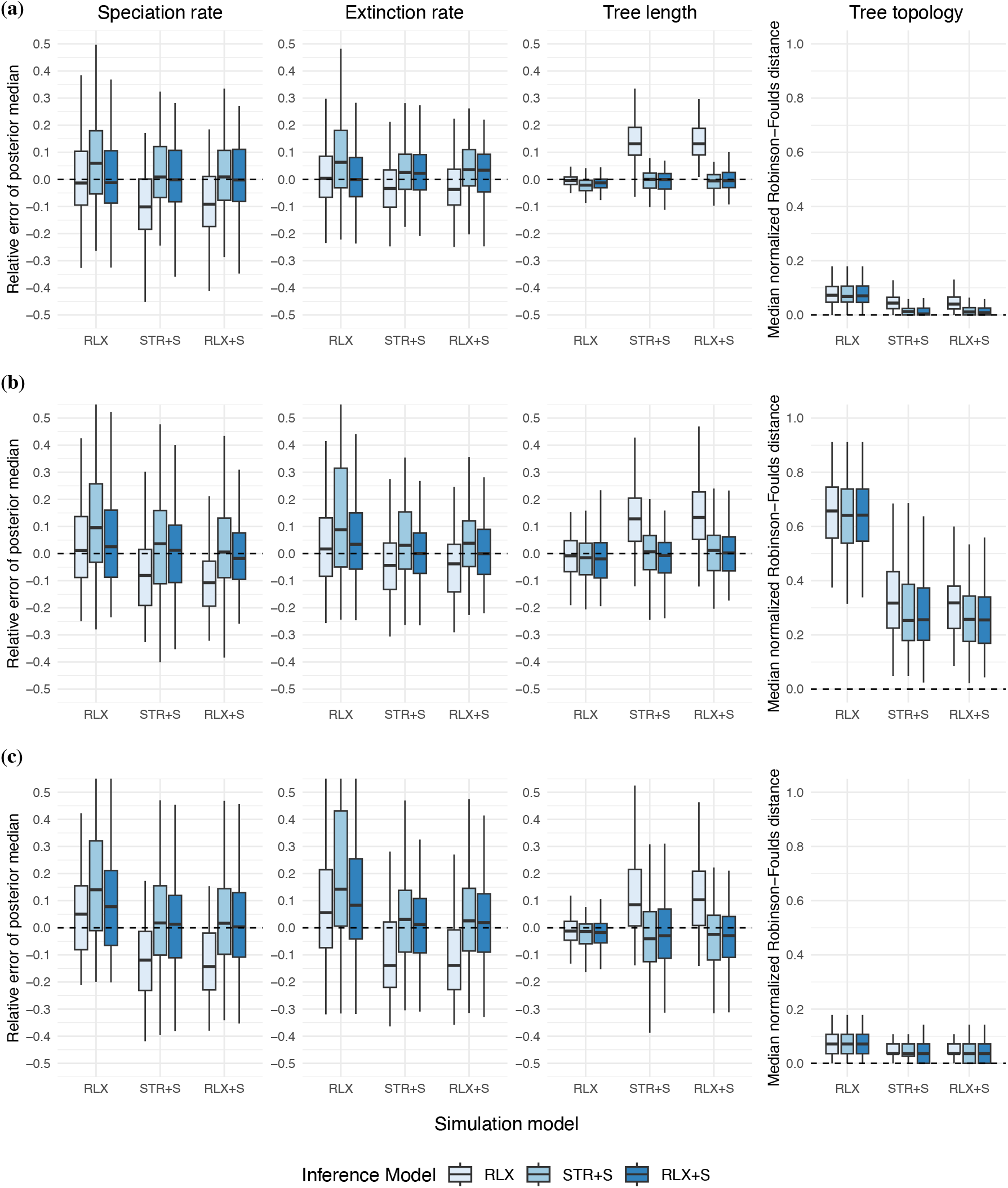
Relative error of posterior median of speciation rate, extinction rate, and total tree length and median normalized Robinson-Foulds distance of posterior trees. Three inference scenarios (rows) are shown: (a) complete sequences, (b) 1% fossil sequences, (c) extant samples only. Each box represents 100 replicates.

The overall estimation accuracy of branching parameters and tree lengths under the correct clock model remained high even when 99% of the sequence data of fossil samples were masked, with median relative errors between −0.02 and 0.04 (Table S12; Figure 4b). The interquantile ranges of relative errors were however wider than those in the complete sequence scenario, indicating increased uncertainty in parameter estimates when the sequence data of fossil samples became sparse. In general, we observed the same patterns of estimation accuracy across the three clock models as in the complete sequence scenario. The RLX+S model remained robust to model misspecification, producing relatively unbiased estimates of branching parameters and tree lengths regardless of the true simulation model (median relative errors between −0.02 and 0.03). Applying the STR+S model to data simulated under RLX led to overestimated speciation rates (∼ 10%) and extinction rates (∼ 9%), whereas the total tree length was relatively unaffected (∼ 2% underestimated), with similar relative errors across all three inference models. When analyzing data simulated under spike clock models, the RLX model produced slightly more biased estimates of speciation rate (∼ 8% − 11% underestimated) and extinction rate (∼ 4% underestimated), while the tree length was overestimated by 13%. Masking 99% of the sequence data for fossil tips drastically increased errors in tree topology estimation, with median normalized Robinson-Foulds distance ranging between 0.25 and 0.66, reflecting the decrease of phylogenetically informative data (Table S15; Figure 4b). Similar to the complete sequence scenario, all three clock models produced similar topological errors when analyzing data simulated without spikes (median distance: 0.64 − 0.66). Using the RLX model to analyze data simulated under spike clock models resulted in an increase in the topological error of the estimated trees (median distance: 0.32) compared to spike clock models (median distance: 0.25 − 0.26), thus showing that even with incomplete data, the choice of clock models can influence the accuracy of tree inference.

When fossil samples were removed from the dataset altogether, the interquantile ranges of relative errors further widened as a result of the reduced data. Nevertheless, the estimation accuracy of branching parameters and tree lengths remained high for spike clock models when they were applied to data simulated with spikes, with median relative errors between −0.04 and 0.03 (Table S13; Figure 4c). While tree length was accurately inferred (median relative error: −0.01), the RLX model tended to overestimate the speciation and extinction rates by 5% − 6% on average when analyzing data simulated under the same model. Compared to the previous two scenarios, the robustness of the RLX+S model decreased when applied to data simulated under RLX, producing estimates of speciation and extinction rates with median relative errors centred around 0.08, whereas applying the STR+S model to data simulated without spikes gave rise to greater overestimation of speciation and extinction rates (∼ 14%). All three clock models gave similarly accurate tree length estimates when analyzing data simulated under RLX, with median relative errors between −0.02 and −0.01. When analyzing data simulated under spike clock models, the RLX model produced more strongly underestimated speciation rates (∼ 12% − 14%) and extinction rates (∼ 14%), while tree length was overestimated by 9% − 10%. All three models estimated highly accurate extant tree topologies regardless of the true simulation model, with median normalized Robinson-Foulds distances between 0.04 and 0.07 (Table S16; Figure 4c). We found no clear increase or decrease in topological error between the three clock models.

## 5 Discussion

Spike clock models were introduced recently as a class of clock models that associate sequence evolution with the branching process of phylogenetic trees. While relaxed clock models are designed to accommodate any and all branch-specific rate variation in a non-parametric fashion, spike models explicitly tie rate variation to the lineage diversification process in that speciation events (observed and inferred) drive character evolution. In this study, we explored the support for spikes across a range of empirical datasets and the robustness of different clock models to model misspecification and missing data.

While our model testing results indicate that no single clock model consistently provides the best fit across all empirical datasets, we find evidence for speciation-associated evolutionary spikes in six of the seven clades analyzed. This widespread signal for spikes suggests that the speciation process can be linked to events of rapid divergence that leave a detectable signature on the alignments of genetic sequences and morphological traits across clades, consistent with the concept of punctuated evolution (Eldredge and Gould 1972; Gould and Eldredge 1977; Katsnelson *et al*. 2019). At macroevolutionary scales, spikes at speciation events can be linked to short phases of rapid evolution or segregated inheritance of instraspecific variance in the parent lineage, for instance, through the emergence of assortative mating or separation of distinct populations (Douglas *et al*. 2025; Duchen *et al*. 2021). While we may not reliably generalize our findings beyond the set of clades analyzed here, the model testing approach we used provides a framework to formally evaluate the prevalence of punctuated evolution across molecular and morphological datasets.

The strict clock model with spikes in which evolutionary rate variation only occurs at speciation events in the form of spikes, receives support as the best-fitting model in three of the clades analyzed, suggesting that spikes alone are sufficient to capture rate heterogeneity in certain empirical datasets. Even in datasets where the STR+S model was not selected as the best-fitting model, its statistical support over the strict clock was comparable to that of relaxed clock models, indicating that speciation-associated spikes are a plausible explanation for the observed rate variation in the empirical data.

Previous analyses of the cephalopod dataset included in our study showed strong evidence for a spike clock model (Douglas *et al*. 2025). While we were able to reproduce the same result, we also found that accounting for incomplete taxon sampling led to different conclusions. Indeed, after accounting for the fact that the extant tips in the tree only represent about 3% of the living cephalopod diversity, a relaxed clock with no spikes received significantly higher support than the alternatives. Earlier research has established that incorrect assumptions about sampling fraction could bias the estimation of speciation and extinction rates (Yang and Rannala 1997; Stadler 2009). Due to the coupling between the speciation process and sequence evolution under spike clock models, this bias appears to propagate into the selection of clock models. The lack of support for spike clock models in the cephalopod dataset does not necessarily mean that evolutionary spikes did not occur in cephalopods, but rather that the alignment data analyzed here do not provide sufficient evidence. Cephalopods have a long evolutionary history of more than 400 million years and extremely high diversity of both living and extinct lineages, which necessarily entails violations of several model assumptions in our analyses. For instance, cephalopods have experienced high levels of variation in speciation and extinction rates both through time and among lineages, having lived through all major Phanerozoic mass extinctions and events of pulsed diversification and selective extinction (Kroger and Yun-Bai 2009; Liu *et al*. 2026). This complex evolutionary history might be inadequately captured by the constant-rate birth-death model applied here, potentially impacting our ability to accurately estimate clock dynamics. Additionally, the sparse sampling and the relatively small morphological character matrix of the analyzed dataset might have obscured the signal of spikes, resulting in the RLX model as a better explanation for the rate variation observed in the data.

In empirical analyses, spike clock models tend to produce more constrained estimates of speciation and extinction rates with lower median values than those inferred under the RLX model, most notably in cephalopods, canids, proboscideans, shrews, and caviomorphs. Compared to RLX, spike clock models inferred speciation and extinction rates that are more consistent with estimates independently derived from the fossil record data in canids and proboscideans (Silvestro *et al*. 2015; Hauffe *et al*. 2026). Indeed, under spike clock models, unsampled lineages are taken into account, as these unobserved speciation events leave behind footprints in the alignment data that are informative of the underlying branching process. We interpret this as the consequence of information transfer from the sequence alignments into the inference of the branching process, as the speciation and extinction rates are estimated jointly with the clock model parameters. In contrast, the topology and branch lengths of estimated trees were relatively consistent across all three clock models, suggesting that the sequence alignments and the age information together provide enough signals to estimate the phylogenetic relationships and divergence times of the sampled taxa, and whether accounting for spikes or not does not significantly influence the overall tree estimates.

Our simulation study demonstrates that the RLX+S model is the most robust to clock model misspecification, giving relatively unbiased and accurate estimates of branching parameters and tree length regardless of the true generating model. While this is expected since both RLX and STR+S are special cases of RLX+S, it also shows that accounting for both spikes and branch rate variation does not lead to obvious overfitting effects. In contrast, both the RLX and STR+S models are prone to biases in parameter estimates under misspecified conditions. Applying the RLX model to data simulated under spike clock models leads to underestimated speciation and extinction rates and overestimated tree lengths. By ignoring spikes in the sequence data, the RLX model attributes all substitution changes to the continuous process of sequence evolution along branches, which results in inflated tree lengths and correspondingly lower speciation and extinction rates. The opposite pattern is observed when the STR+S model is applied to data that did not evolve under a spike model, which leads to overestimation of speciation and extinction rates, although the tree length estimates remain relatively accurate. As rate variation is modeled exclusively through spikes, the STR+S model seemingly compensates by overestimating speciation and extinction rates to introduce more unobserved speciation events in the branching process and, in turn, more spikes to explain the rate patterns generated by the RLX model.

As we degrade fossil sequence data and eventually remove fossil samples altogether, the overall parameter estimation error increases, and biases under the misspecified condition become stronger. Even when analyzing data simulated under the correct model, the RLX model begins to produce less accurate estimates of branching parameters (especially extinction rate) as fewer data are used. In contrast, spike clock models are robust to missing data, producing accurate estimates of branching parameters even when fossil tips are completely removed from the sample. This result indicates that evolutionary spikes are informative of the underlying branching process, and accounting for them in the clock model can help mitigate the effect of missing data on branching parameter estimation.

The simulation study did not recover the same pattern observed in the empirical analyses, namely that spike clock models give lower median estimates and more constrained uncertainty intervals of speciation and extinction rates compared to RLX in some datasets. When applied to simulated data, spike clock models tend to produce higher speciation and extinction rates than the RLX model under all simulation conditions. This disparity might be attributed to the differences between the simulated data and the real-world observations which are generated by much more complex and heterogeneous evolutionary processes. All clock models are misspecified when applied to empirical data for which we do not know the true generating processes (Ducĥene *et al*. 2015). Moreover, most empirical datasets include morphological characters for fossil samples, a data type not investigated in our simulations and for which adequate evolutionary modeling may still be difficult to achieve (Mulvey *et al*. 2025).

## 6 Conclusions

As clock models are central to our understanding of the diversification of clades, methodological advances linking molecular and morphological evolution with the speciation process offer new opportunities to achieve more mechanistic modeling of rate variation. In our study we compared different clock models and found that evolutionary spikes receive consistent support in empirical datasets. Our analyses also showed that even a strict clock when coupled with spikes can receive strong model support, suggesting that it is able to capture a substantial fraction of the empirical signal for rate variation.

The relaxed clock model with spikes is a robust model choice, as it produces relatively accurate estimates of branching parameters and tree lengths, even when the true underlying process does not involve spikes. Yet, correctly accounting for spikes is beneficial for accurate estimation of tree topologies and branching parameters. This becomes even more important when the sequence data of fossil samples are sparse or completely missing, because spikes leave signals in the sampled sequence data that are informative of the underlying branching process and phylogenetic relationships of sampled taxa.

Current clock models with spikes likely remain far too simple to fully capture the complex dynamics of evolution. For instance, the assumption of a constant-rate birth-death process is invariably violated in empirical systems, and the impact of these model violations on clock model selection and parameter estimation under spike clock models remains unclear. Future developments should consider more complex branching processes under spike clock models, such as time- or lineage-dependent variation in diversification rates. This additional complexity should lead to more mechanistic and interpretable models that better capture the complex nature of macroevolutionary processes.

## 7 Code and data availability

All code and data associated with this paper are available in the project repository. The source code for the Multi-type Spike Model is available in the Multi-type Spike Model repository.

## 9 Acknowledgments

The authors would like to thank Ugnė Stolz and Tobias Dieselhorst for helpful discussions and feedback.

## 10 Supplementary Information

### 10.1 Summary of clock models

Under a strict clock (STR) model, the rate of evolution is constant across all branches of a phylogeny:

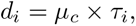

where the branch length *d_i_* represents the genetic or morphological distance for a branch *i*, measured in the expected number of substitutions per site. *d_i_* is the product of the time duration of that branch *τ_i_* and the mean clock rate *µ_c_*, measured in the expected number of substitutions per site per unit time.

Under a relaxed clock (RLX) model, each branch is assigned a unique clock rate by the relative rate modifier *r_i_*:

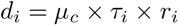

The relative rates are independently drawn from a log-normal distribution whose variation is controlled by a standard deviation parameter *σ_r_*:

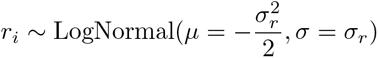

Under a strict clock model with spikes (STR+S), while the clock rate is constant across all branches, evolutionary rate variation occurs through spikes at speciation events:

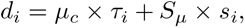

where the mean spike size (*S_µ_*) describes the expected number of sudden substitutions per site per speciation, and each branch is assigned a unique spike size by the relative spike size modifier *s_i_*. The relative spike sizes are drawn from a gamma distribution whose variation is controlled by a shape parameter *S_α_*:

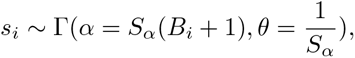

where *B_i_* is the number of unobserved speciation events on branch *i*. The model takes into account both observed and unsampled speciation events to estimate the relative spike size on each branch.

Under a relaxed clock model with spikes (RLX+S), both clock rates and spike sizes vary between branches:

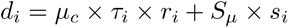

Under spike clock models (*i.e.*, STR+S and RLX+S), we integrate over the number of unobserved speciation events *B_i_*and calculate the marginal probability of relative spike size *s_i_* for each branch *i*:

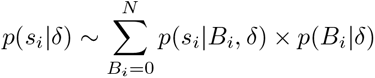

for some large *N* such that the cumulative Poisson probability Σ*_Bi_ p*(*B_i_*|*δ*) *>* 0.99, where *δ* = (*λ, µ, ψ, ρ*) is the vector of branching parameters.

The probability of relative spike size along a branch with *B_i_* unobserved speciation events is

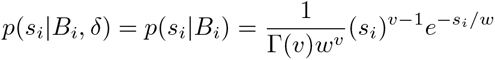

where *v* = *S_α_*(*B_i_* + 1) and *w* = 1*/S_α_*.

The probability of having *B_i_* unobserved speciation events on a branch with interval [*t_o_, t_e_*] follows a Poisson distribution:

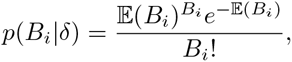

where the expected number of unobserved speciation events on a branch with interval [*t_o_, t_e_*], E(*B_i_*), can be evaluated by

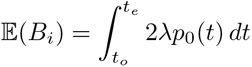

following Stolz *et al*. (2024).

### 10.2 Inference model set-up of empirical analyses

#### 10.2.1 Cephalopoda

We adapted the XML file from Douglas *et al*. (2025) to set up the inference model for cephalopods. The original data came from Whalen and Landman (2022).

**Substitution model:** Morphological data were divided into five partitions based on the number of states (two to six states). We applied the Mk model (Lewis 2001) with ascertainment correction and gamma-distributed across-site rate variation with four categories. The gamma shape parameter was shared across all partitions with an exponential prior Exp(1). We allowed for partition-specific relative substitution rates and fixed the average rate per site to one.

**Clock model:** We used the following priors for the clock model: mean clock rate *µ_c_*∼ LogNormal(mean = 0.0001*, σ* = 2), standard deviation of relative clock rates *σ_r_* ∼ Γ(*α* = 5*, θ* = 0.05), mean spike size *S_µ_*∼ LogNormal(mean = 0.01*, σ* = 1.2), shape parameter of relative spike sizes *S_α_* ∼ LogNormal(mean = 2*, σ* = 0.5).

**Tree model:** We used the fossilized birth-death model with the following priors: speciation rate *λ* ∼ LogNormal(mean = 0.001*, σ* = 2), reproductive number *r*_0_ ∼ LogNormal(mean = 0.2*, σ* = 1) with an offset set to one, sampling proportion 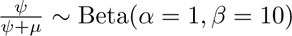, origin time ∼ U(0, 4600), extant sampling fraction *ρ* = 1 or *ρ* = 0.03375.

**Age calibration:** All fossil samples were included as tips whose ages were fixed to single point estimates without ranges. The ages of all extant samples were fixed to zero. More detailed age information is included in the provided XML file in the Supplementary Materials.

#### 10.2.2 Canidae

We adapted the XML file from Stolz *et al*. (2025) to set up the inference model for canids. The original data came from Slater (2015).

**Substitution model:** Morphological data were divided into four partitions based on the number of states (two to five states). We applied the Mk model (Lewis 2001) without ascertainment correction or across-site rate variation. All partitions evolved at the same substitution rate.

**Clock model:** We used the following priors for the clock model: mean clock rate *µ_c_*∼ Exp(mean = 1), standard deviation of relative clock rates *σ_r_* ∼ Γ(*α* = 5*, θ* = 0.05), mean spike size *S_µ_*∼ LogNormal(mean = 0.01*, σ* = 1.2), shape parameter of relative spike sizes *S_α_* ∼ LogNormal(mean = 2*, σ* = 0.5).

**Tree model:** We used the fossilized birth-death model with the following priors: net diversification rate *λ* − *µ* ∼ Exp(mean = 1), turnover rate 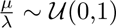, sampling proportion 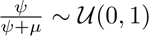, origin time ∼ U(0, Inf), extant sampling fraction *ρ* ∼ U(0, 1). The empirical sampling fraction of extant canids is 5*/*35 = 0.143 based on the species count by Wilson and Reeder (2005).

**Age calibration:** All fossil samples were included as tips whose ages were uniformly sampled between the oldest estimate of the first occurrence and the youngest estimate of the last occurrence. The ages of all extant samples were fixed to zero. More detailed age information is included in the provided XML file in the Supplementary Materials.

#### 10.2.3 Platyrrhini

We adapted the XML file from Silvestro *et al*. (2019) to set up the inference model for platyrrhines.

**Substitution model:** DNA data were divided into seven partitions based on chromosomal positions (four autosomal, two allosomal, and one mitochondrial). For each partition, we applied the GTR model (Tavaŕe 1986) with estimated equilibrium base frequencies and gamma-distributed across-site rate variation with four categories. We used Γ(*α* = 0.05*, θ*= 10) for the transversion rates and Γ(*α* = 0.05*, θ* = 20) for the transition rates of the GTR model, U(0, 1) for the equilibrium base frequencies, and the gamma shape parameter fixed to one. We allowed for partition-specific relative substitution rates and fixed the average rate per site to one.

**Clock model:** We used the following priors for the clock model: mean clock rate *µ_c_*∼ Exp(mean = 10), standard deviation of relative clock rates *σ_r_* ∼ Exp(mean = 1), mean spike size *S_µ_* ∼ LogNormal(mean = 0.01*, σ* = 1.2), shape parameter of relative spike sizes *S_α_* ∼ LogNormal(mean = 2*, σ* = 0.5).

**Tree model:** We used the fossilized birth-death model with the following priors: net diversification rate *λ* − *µ* ∼ Exp(mean = 1), turnover rate 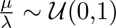, sampling proportion 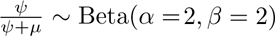, origin time ∼ Γ(*α* = 2*, θ* = 5) with an offset set to 37.5, extant sampling fraction *ρ* = 0.68.

**Age calibration:** All fossil samples were included as tips whose ages were fixed to single point estimates without ranges. The ages of all extant samples were fixed to zero. We employed 20 monophyletic topological constraints to enforce the phylogenetic placement of the fossils. More detailed age information is included in the provided XML file in the Supplementary Materials.

#### 10.2.4 Proboscidea

We adapted the XML file from Baleka *et al*. (2022) to set up the inference model for proboscideans.

**Substitution model:** All DNA data were placed into a single partition (mitochondrial). We applied the HKY model (Hasegawa *et al*. 1985) with estimated equilibrium base frequencies and gamma-distributed across-site rate variation with four categories. We used LogNormal(*µ* = 1*, σ* = 1.25) for the ratio of transition rates *versus* transversion rates of the HKY model, U(0, 1) for the equilibrium base frequencies, and Exp(mean = 1) for the gamma shape parameter. Morphological characters were divided into five partitions based on the number of states (two to six states). We applied the Mk model (Lewis 2001) with ascertainment correction and gamma-distributed across-site rate variation with four categories. The gamma shape parameter was shared across all morphological partitions with an exponential prior Exp(mean = 1). All morphological partitions evolved at the same substitution rate.

**Clock model:** We applied two separate clocks for the two data types with the following priors: mean clock rate *µ_c_* ∼ Exp(mean = 1), standard deviation of relative clock rates *σ_r_* ∼ Γ(*α* = 5*, θ* = 0.05), mean spike size *S_µ_* ∼ LogNormal(mean = 0.01*, σ* = 1.2), shape parameter of relative spike sizes *S_α_* ∼ LogNormal(mean = 2*, σ* = 0.5).

**Tree model:** We used the fossilized birth-death model with the following priors: speciation rate *λ* ∼ U(0, 100), extinction rate *µ* ∼ U(0,100), sampling rate *ψ* ∼ U(0, 10), origin time ∼ U(0, 4600), extant sampling fraction *ρ* = 1.

**Age calibration:** All fossil samples were included as tips whose ages were uniformly sampled between the oldest and the youngest boundaries of their stratigraphic ranges. The ages of all extant samples were fixed to zero. We applied two monophyletic topological contraints and four node age calibrations with normal distributions. More detailed age information is included in the provided XML file in the Supplementary Materials.

#### 10.2.5 Spheniscidae

We adapted the XML file from Gavryushkina *et al*. (2017) to set up the inference model for penguins.

**Substitution model:** DNA data were divided into five partitions based on locus positions (four mitochondrial loci and one nuclear gene). For each partition, we applied the GTR model (Tavaŕe 1986) with estimated equilibrium base frequencies and gamma-distributed across-site rate variation with four categories. We used U(0, 1) for the substitution rates and equilibrium base frequencies of the GTR model and U(0, 10) for the gamma shape parameter. We allowed for partition-specific relative substitution rates and fixed the average rate per site to one. Morphological data were divided into six partitions based on the number of states (two to seven states). We applied the Mk model (Lewis 2001) with ascertainment correction and gamma-distributed across-site rate variation with four categories. The gamma shape parameter was shared across all morphological partitions with a uniform prior U(0, 10). All morphological partitions evolved at the same substitution rate.

**Clock model:** We applied two separate clocks for the two data types with the following priors: mean clock rate *µ_c_* ∼ LogNormal(*µ* = −3.5*, σ* = 1) for DNA data and LogNormal(*µ* = −5.5*, σ* = 2) for morphological characters, standard deviation of relative clock rates *σ_r_* ∼ Γ(*α* = 0.5396*, θ* = 0.3819), mean spike size *S_µ_* ∼ LogNormal(mean = 0.01*, σ* = 1.2), shape parameter of relative spike sizes *S_α_* ∼ LogNormal(mean = 2*, σ* = 0.5).

**Tree model:** We used the fossilized birth-death model with the following priors: net diversification rate *λ* − *µ* ∼ LogNormal(*µ* = −3.5*, σ* = 1), turnover rate 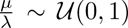, sampling proportion 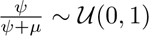 U(0, 1), origin time ∼ U(0, 160), extant sampling fraction *ρ* = 1.

**Age calibration:** All fossil samples were included as tips whose ages were uniformly sampled between the oldest and the youngest boundaries of their stratigraphic ranges. The ages of all extant samples were fixed to zero. More detailed age information is included in the provided XML file in the Supplementary Materials.

#### 10.2.6 Soricidae

We adapted the XML file from Yuan *et al*. (2024) to set up the inference model for shrews.

**Substitution model:** DNA data were divided into three partitions: nuclear genes, mitochondrial genes, and ribosomal DNA. For each partition, we applied the GTR model (Tavaŕe 1986) with empirical equilibrium base frequencies and gamma-distributed across-site rate variation with four categories. We used Γ(*α* = 0.05*, θ* = 10) for the transversion rates and Γ(*α* = 0.05*, θ* = 20) for the transition rates of the GTR model and Exp(mean = 1) for the shape parameter of the gamma-distributed rate variation. We allowed for partition-specific relative substitution rates and fixed the average rate per site to one. Morphological data were divided into five partitions based on the number of states (two to six states). We applied the Mk model (Lewis 2001) with ascertainment correction and gamma-distributed across-site rate variation with four categories. The gamma shape parameter was independent for each morphological partition with an exponential prior Exp(mean = 1). All morphological partitions evolved at the same substitution rate.

**Clock model:** We applied two separate clocks for the two data types with the following priors: mean clock rate *µ_c_* ∼ Γ(*α* = 2.1*, θ* = 115.52), standard deviation of relative clock rates *σ_r_* ∼ Γ(*α* = 5*, θ* = 0.05), mean spike size *S_µ_*∼ LogNormal(mean = 0.01*, σ* = 1.2), shape parameter of relative spike sizes *S_α_* ∼ LogNormal(mean = 2*, σ* = 0.5).

**Tree model:** We used the fossilized birth-death model with the following priors: speciation rate *λ* ∼ Exp(mean = 10), extinction rate *µ* ∼ Beta(*α* = 1*, β* = 1), sampling rate *ψ* ∼ Beta(*α* = 1*, β* = 1), root age ∼ Γ(*α* = 1.3*, θ* = 20) with an offset set to 61.66, extant sampling fraction *ρ* = 0.09.

**Age calibration:** All fossil samples were included as tips whose ages were uniformly sampled between the oldest and the youngest boundaries of their stratigraphic ranges. The ages of all extant samples were fixed to zero. More detailed age information is included in the provided XML file in the Supplementary Materials.

#### 10.2.7 Caviomorpha

We adapted the XML file from Carrillo *et al*. (2026) to set up the inference model for caviomorphs.

**Substitution model:** All DNA data were placed into a single partition. We applied the GTR model (Tavaŕe 1986) with estimated equilibrium base frequencies and gamma-distributed across-site rate variation with four categories. We used Γ(*α* = 0.05*, θ* = 10) for the transversion rates and Γ(*α* = 0.05*, θ* = 20) for the transition rates of the GTR model, U(0, 1) for the equilibrium base frequencies, and Exp(mean = 1) for the shape parameter of the gamma-distributed rate variation. Morphological data were divided into five partitions based on the number of states (two to six states). We applied the Mk model (Lewis 2001) without ascertainment correction, while allowing for gamma-distributed rate variation with four categories. The gamma shape parameter was independent for each morphological partition with an exponential prior Exp(mean = 1). All morphological partitions evolved at the same substitution rate.

**Clock model:** We applied two separate clocks for the two data types with the following priors: mean clock rate *µ_c_* ∼ Exp(mean = 1), standard deviation of relative clock rates *σ_r_* ∼ Exp(mean = 1), mean spike size *S_µ_* ∼ LogNormal(mean = 0.01*, σ* = 1.2), shape parameter of relative spike sizes *S_α_*∼ LogNormal(mean = 2*, σ* = 0.5).

**Tree model:** We used the fossilized birth-death model with the following priors: net diversification rate *λ* − *µ* ∼ Exp(mean = 1), turnover rate 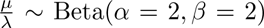, sampling proportion 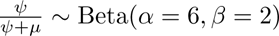, origin time fixed to 65, extant sampling fraction *ρ* = 1.

**Age calibration:** All fossil samples were included as tips whose ages were uniformly sampled between the oldest and the youngest boundaries of their stratigraphic ranges. The ages of all extant samples were fixed to zero. We applied 18 monophyletic topological constraints. More detailed age information is included in the provided XML file in the Supplementary Materials.

**Table S1:** Summary of seven empirical datasets investigated in this study. The table includes the clade name, data source, number of extant and fossil samples, number of DNA sites and morphological traits, and the approximate geological period of the root of the clade.

| Clade | Source | Extant taxa | $\rho$ | Fossils | DNA sites | Traits | Root age |
| --- | --- | --- | --- | --- | --- | --- | --- |
| Cephalopoda | Whalen and Landman<br>(2022) | 27 | 0.03375 | 52 | 0 | 153 | Cambrian |
| Canidae | Slater (2015) | 5 | 0.143 | 116 | 0 | 123 | Eocene |
| Platyrrhini | Silvestro *et al.*<br>(2019) | 87 | 0.68 | 34 | 36065 | 0 | Eocene |
| Proboscidea | Baleka *et al.*<br>(2022) | 3 | 1 | 33 | 2929 | 125 | Eocene |
| Spheniscidae | Gavryushkina<br>*et al.* (2017) | 19 | 1 | 36 | 8145 | 202 | Eocene |
| Soricidae | Yuan *et al.*<br>(2024) | 44 | 0.09 | 23 | 18762 | 217 | Paleocene |
| Caviomorpha | Carrillo *et al.*<br>(2026) | 149 | 1 | 52 | 33850 | 513 | Oligocene |

**Table S2:** Marginal likelihood estimates of seven empirical datasets under four clock models. The table includes the number of replicates and steps for each model, the mean log marginal likelihood estimate (*logZ*^^^), and the standard deviation (SD) of the estimates across replicates.

| Clade | Model | # replicates | # steps | $\log \hat{Z}$ | SD |
| --- | --- | --- | --- | --- | --- |
| Cephalopoda ($\rho = 0.03375$) | STR | 10 | 30 | -2442.5 | 0.203769 |
|  | RLX | 10 | 30 | -2429 | 0.210171 |
|  | STR+S | 10 | 30 | -2435.84 | 1.56352 |
|  | RLX+S | 10 | 30 | -2436.58 | 1.61369 |
| Cephalopoda ($\rho = 1$) | STR | 10 | 30 | -2450.22 | 0.456585 |
|  | RLX | 10 | 30 | -2435.17 | 0.682522 |
|  | STR+S | 10 | 30 | -2424.12 | 1.3428 |
|  | RLX+S | 10 | 30 | -2424.18 | 1.15292 |
| Canidae | STR | 10 | 30 | -3242.21 | 0.518799 |
|  | RLX | 10 | 30 | -3219.35 | 1.06341 |
|  | STR+S | 10 | 30 | -3208.76 | 1.35477 |
|  | RLX+S | 10 | 30 | -3208.59 | 2.27256 |
| Platyrrhini | STR | 10 | 60 | -119436 | 0.921213 |
|  | RLX | 10 | 60 | -119326 | 1.27423 |
|  | STR+S | 10 | 60 | -119319 | 0.829641 |
|  | RLX+S | 10 | 60 | -119325 | 0.808196 |
| Proboscidea | STR | 10 | 30 | -6109.82 | 0.0523788 |
|  | RLX | 10 | 30 | -6087.01 | 0.0760294 |
|  | STR+S | 10 | 30 | -6082.04 | 0.44659 |
|  | RLX+S | 10 | 30 | -6081.81 | 0.990548 |
| Spheniscidae | STR | 10 | 30 | -19703.1 | 0.418161 |
|  | RLX | 10 | 30 | -19678.9 | 0.838246 |
|  | STR+S | 10 | 30 | -19680.4 | 0.932574 |
|  | RLX+S | 10 | 30 | -19675.9 | 1.18321 |
| Soricidae | STR | 10 | 60 | -226116 | 0.987415 |
|  | RLX | 10 | 60 | -225842 | 0.977142 |
|  | STR+S | 10 | 60 | -225851 | 0.986515 |
|  | RLX+S | 10 | 60 | -225838 | 1.18686 |
| Caviomorpha | STR | 10 | 60 | -164252 | 1.11509 |
|  | RLX | 10 | 60 | -163821 | 1.28516 |
|  | STR+S | 10 | 60 | -163828 | 1.48498 |
|  | RLX+S | 10 | 60 | -163802 | 1.4928 |

**Table S3:** Posterior estimates of speciation rate, extinction rate, and total branch length for Cephalopoda. ESS = effective sample size, N = number of posterior samples.

| Model | Parameter | Mean | Median | 95% HPD lower | 95% HPD upper | ESS | N |
| --- | --- | --- | --- | --- | --- | --- | --- |
| RLX | Speciation rate | 0.392 | 0.372 | 0.185 | 0.656 | 1400.000 | 1400 |
|  | Extinction rate | 0.385 | 0.366 | 0.182 | 0.654 | 1400.000 | 1400 |
|  | Tree length | 4990.080 | 4943.561 | 4030.376 | 6016.467 | 855.798 | 1400 |
| RLX+S | Speciation rate | 0.165 | 0.158 | 0.077 | 0.265 | 767.253 | 3762 |
|  | Extinction rate | 0.159 | 0.152 | 0.068 | 0.255 | 760.386 | 3762 |
|  | Tree length | 5731.177 | 5650.620 | 4495.786 | 7151.753 | 1647.403 | 3762 |
| STR+S | Speciation rate | 0.169 | 0.162 | 0.080 | 0.266 | 556.510 | 3222 |
|  | Extinction rate | 0.162 | 0.156 | 0.073 | 0.259 | 555.024 | 3222 |
|  | Tree length | 5682.457 | 5630.283 | 4413.230 | 6945.574 | 1194.951 | 3222 |

**Table S4:** Posterior estimates of speciation rate, extinction rate, and total branch length for Canidae (dogs). ESS = effective sample size, N = number of posterior samples.

| Model | Parameter | Mean | Median | 95% HPD lower | 95% HPD upper | ESS | N |
| --- | --- | --- | --- | --- | --- | --- | --- |
| RLX | Speciation rate | 1.039 | 0.958 | 0.488 | 1.791 | 26628.098 | 45005 |
|  | Extinction rate | 0.972 | 0.890 | 0.413 | 1.703 | 25575.505 | 45005 |
|  | Tree length | 267.661 | 266.693 | 235.709 | 301.573 | 6604.958 | 45005 |
| RLX+S | Speciation rate | 0.783 | 0.768 | 0.485 | 1.088 | 312.728 | 10995 |
|  | Extinction rate | 0.717 | 0.701 | 0.434 | 1.026 | 308.372 | 10995 |
|  | Tree length | 263.831 | 263.020 | 232.929 | 296.849 | 987.200 | 10995 |
| STR+S | Speciation rate | 0.784 | 0.769 | 0.486 | 1.096 | 1095.966 | 10935 |
|  | Extinction rate | 0.717 | 0.703 | 0.426 | 1.020 | 1066.675 | 10935 |
|  | Tree length | 263.927 | 263.188 | 232.155 | 296.473 | 1924.468 | 10935 |

**Table S5:** Posterior estimates of speciation rate, extinction rate, and total branch length for Platyrrhini (New World monkeys). ESS = effective sample size, N = number of posterior samples.

| Model | Parameter | Mean | Median | 95% HPD lower | 95% HPD upper | ESS | N |
| --- | --- | --- | --- | --- | --- | --- | --- |
| RLX | Speciation rate | 0.360 | 0.355 | 0.251 | 0.498 | 1260.128 | 1948 |
|  | Extinction rate | 0.268 | 0.261 | 0.135 | 0.410 | 1301.892 | 1948 |
|  | Tree length | 645.349 | 639.472 | 552.839 | 745.639 | 553.912 | 1948 |
| RLX+S | Speciation rate | 0.379 | 0.374 | 0.267 | 0.493 | 572.649 | 2638 |
|  | Extinction rate | 0.290 | 0.285 | 0.175 | 0.419 | 528.873 | 2638 |
|  | Tree length | 633.528 | 629.799 | 544.500 | 724.681 | 264.582 | 2638 |
| STR+S | Speciation rate | 0.378 | 0.373 | 0.270 | 0.500 | 467.719 | 2289 |
|  | Extinction rate | 0.288 | 0.284 | 0.166 | 0.411 | 438.375 | 2289 |
|  | Tree length | 631.935 | 626.916 | 545.328 | 724.408 | 462.034 | 2289 |

**Table S6:** Posterior estimates of speciation rate, extinction rate, and total branch length for Proboscidea (elephants and relatives). ESS = effective sample size, N = number of posterior samples.

| Model | Parameter | Mean | Median | 95% HPD lower | 95% HPD upper | ESS | N |
| --- | --- | --- | --- | --- | --- | --- | --- |
| RLX | Speciation rate | 0.981 | 0.811 | 0.227 | 2.257 | 1658.957 | 5005 |
|  | Extinction rate | 0.983 | 0.812 | 0.243 | 2.303 | 1665.777 | 5005 |
|  | Tree length | 176.367 | 175.294 | 144.459 | 208.520 | 3017.103 | 5005 |
| RLX+S | Speciation rate | 0.361 | 0.351 | 0.194 | 0.543 | 2260.899 | 18005 |
|  | Extinction rate | 0.359 | 0.348 | 0.197 | 0.546 | 2286.832 | 18005 |
|  | Tree length | 168.831 | 167.602 | 137.153 | 202.112 | 1833.020 | 18005 |
| STR+S | Speciation rate | 0.374 | 0.363 | 0.204 | 0.570 | 482.628 | 5005 |
|  | Extinction rate | 0.371 | 0.359 | 0.197 | 0.569 | 493.073 | 5005 |
|  | Tree length | 166.867 | 165.325 | 134.382 | 200.942 | 688.697 | 5005 |

**Table S7:** Posterior estimates of speciation rate, extinction rate, and total branch length for Spheniscidae (penguins). ESS = effective sample size, N = number of posterior samples.

| Model | Parameter | Mean | Median | 95% HPD lower | 95% HPD upper | ESS | N |
| --- | --- | --- | --- | --- | --- | --- | --- |
| RLX | Speciation rate | 0.328 | 0.312 | 0.147 | 0.530 | 6609.932 | 7030 |
|  | Extinction rate | 0.278 | 0.262 | 0.102 | 0.495 | 6568.816 | 7030 |
|  | Tree length | 196.271 | 193.600 | 157.119 | 238.043 | 1673.353 | 7030 |
| RLX+S | Speciation rate | 0.346 | 0.331 | 0.172 | 0.568 | 467.738 | 2575 |
|  | Extinction rate | 0.298 | 0.285 | 0.111 | 0.522 | 422.806 | 2575 |
|  | Tree length | 202.130 | 200.972 | 159.644 | 248.300 | 344.729 | 2575 |
| STR+S | Speciation rate | 0.364 | 0.351 | 0.200 | 0.550 | 1109.878 | 12157 |
|  | Extinction rate | 0.318 | 0.304 | 0.151 | 0.500 | 883.634 | 12157 |
|  | Tree length | 206.026 | 204.534 | 172.374 | 241.834 | 1234.622 | 12157 |

**Table S8:** Posterior estimates of speciation rate, extinction rate, and total branch length for Soricidae (shrews). ESS = effective sample size, N = number of posterior samples.

| Model | Parameter | Mean | Median | 95% HPD lower | 95% HPD upper | ESS | N |
| --- | --- | --- | --- | --- | --- | --- | --- |
| RLX | Speciation rate | 0.366 | 0.351 | 0.177 | 0.577 | 3457.591 | 7280 |
|  | Extinction rate | 0.313 | 0.297 | 0.113 | 0.531 | 3567.606 | 7280 |
|  | Tree length | 1297.542 | 1279.578 | 1083.444 | 1527.700 | 1962.736 | 7280 |
| RLX+S | Speciation rate | 0.281 | 0.276 | 0.159 | 0.416 | 421.437 | 7323 |
|  | Extinction rate | 0.221 | 0.215 | 0.096 | 0.365 | 392.514 | 7323 |
|  | Tree length | 1308.290 | 1287.245 | 1091.039 | 1575.088 | 1359.350 | 7323 |
| STR+S | Speciation rate | 0.226 | 0.222 | 0.142 | 0.320 | 345.961 | 14056 |
|  | Extinction rate | 0.163 | 0.159 | 0.075 | 0.254 | 326.066 | 14056 |
|  | Tree length | 1270.915 | 1251.872 | 1066.195 | 1509.169 | 2643.873 | 14056 |

**Table S9:** Posterior estimates of speciation rate, extinction rate, and total branch length for Caviomorpha. ESS = effective sample size, N = number of posterior samples.

| Model | Parameter | Mean | Median | 95% HPD lower | 95% HPD upper | ESS | N |
| --- | --- | --- | --- | --- | --- | --- | --- |
| RLX | Speciation rate | 0.219 | 0.217 | 0.166 | 0.274 | 707.600 | 7977 |
|  | Extinction rate | 0.170 | 0.168 | 0.118 | 0.229 | 975.550 | 7977 |
|  | Tree length | 1603.641 | 1601.646 | 1413.142 | 1796.515 | 303.920 | 7977 |
| RLX+S | Speciation rate | 0.204 | 0.203 | 0.164 | 0.250 | 1263.456 | 10055 |
|  | Extinction rate | 0.152 | 0.151 | 0.109 | 0.195 | 1474.292 | 10055 |
|  | Tree length | 1584.723 | 1582.253 | 1417.472 | 1758.404 | 504.195 | 10055 |
| STR+S | Speciation rate | 0.200 | 0.199 | 0.157 | 0.240 | 455.571 | 4122 |
|  | Extinction rate | 0.149 | 0.147 | 0.108 | 0.195 | 431.193 | 4122 |
|  | Tree length | 1641.851 | 1636.619 | 1479.944 | 1821.802 | 494.600 | 4122 |

**Figure S1:**
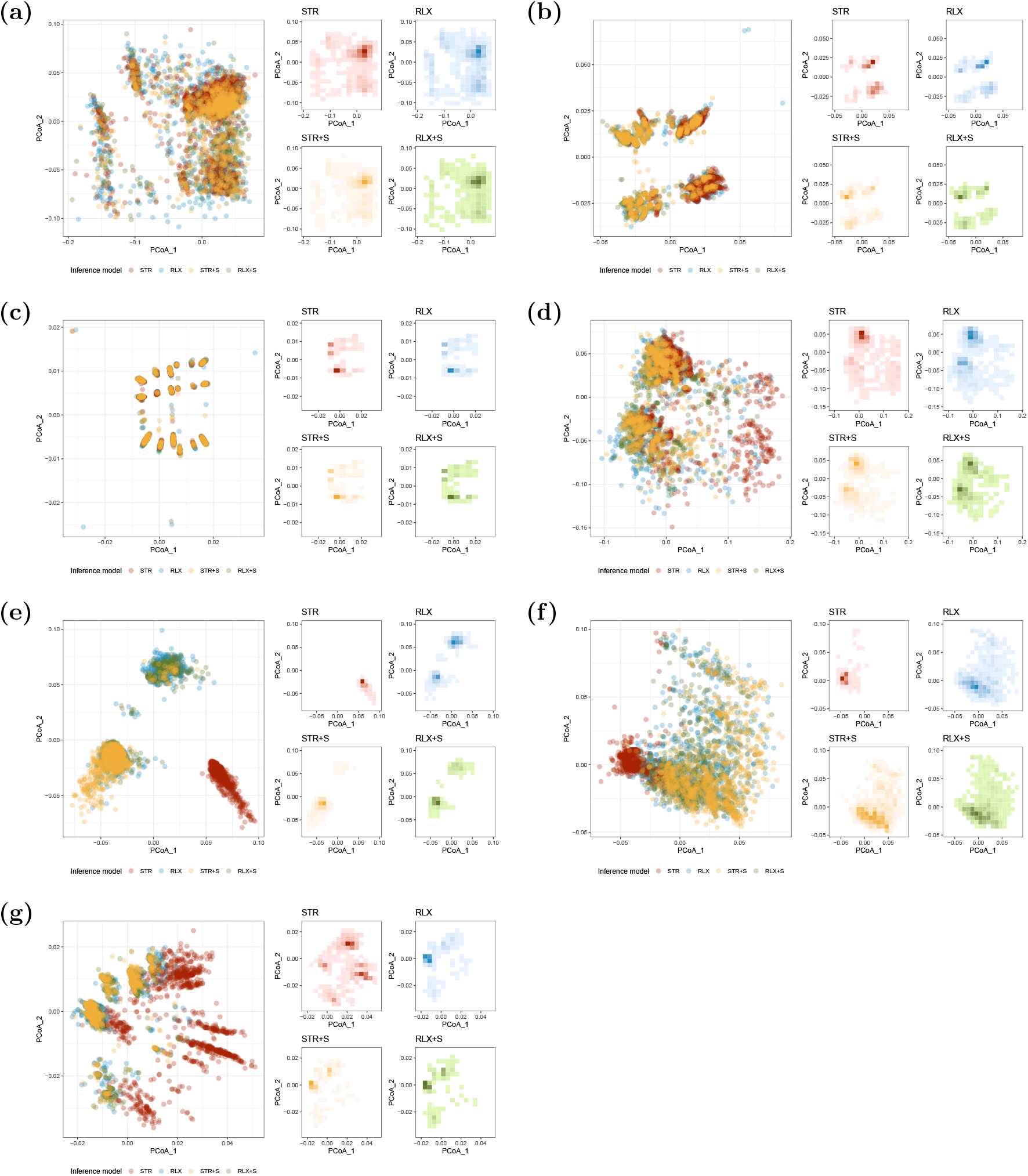
Quartet distance treespace for empirical datasets analysed under four clock models. (a) Cephalopoda with empirical *ρ* sampling probability. (b) Canidae. (c) Platyrrhini. (d) Proboscidea. (e) Spheniscidae. (f) Soricidae. (g) Caviomorpha.

**Table S10:** Summary statistics of 100 simulated trees.

| **Variable** | **Mean** | **Median** | **95% range** | **Min-max range** |
| --- | --- | --- | --- | --- |
| Number of tips | 66.13 | 57 | 36 – 129 | 35 – 137 |
| Number of sampled ancestors | 26.85 | 20 | 6 – 78.15 | 4 – 86 |
| Age of the oldest sample | 13.86 | 9.40 | 1.90 – 37.21 | 1.43 – 45.45 |
| Origin time | 16.79 | 12.30 | 4.09 – 42.73 | 2.71 – 50.31 |
| Root age | 15.34 | 11.03 | 3.07 – 38.99 | 2.41 – 46.77 |
| Total branch length | 129.43 | 99.88 | 30.43 – 369.56 | 27.48 – 390.29 |

**Table S11:**
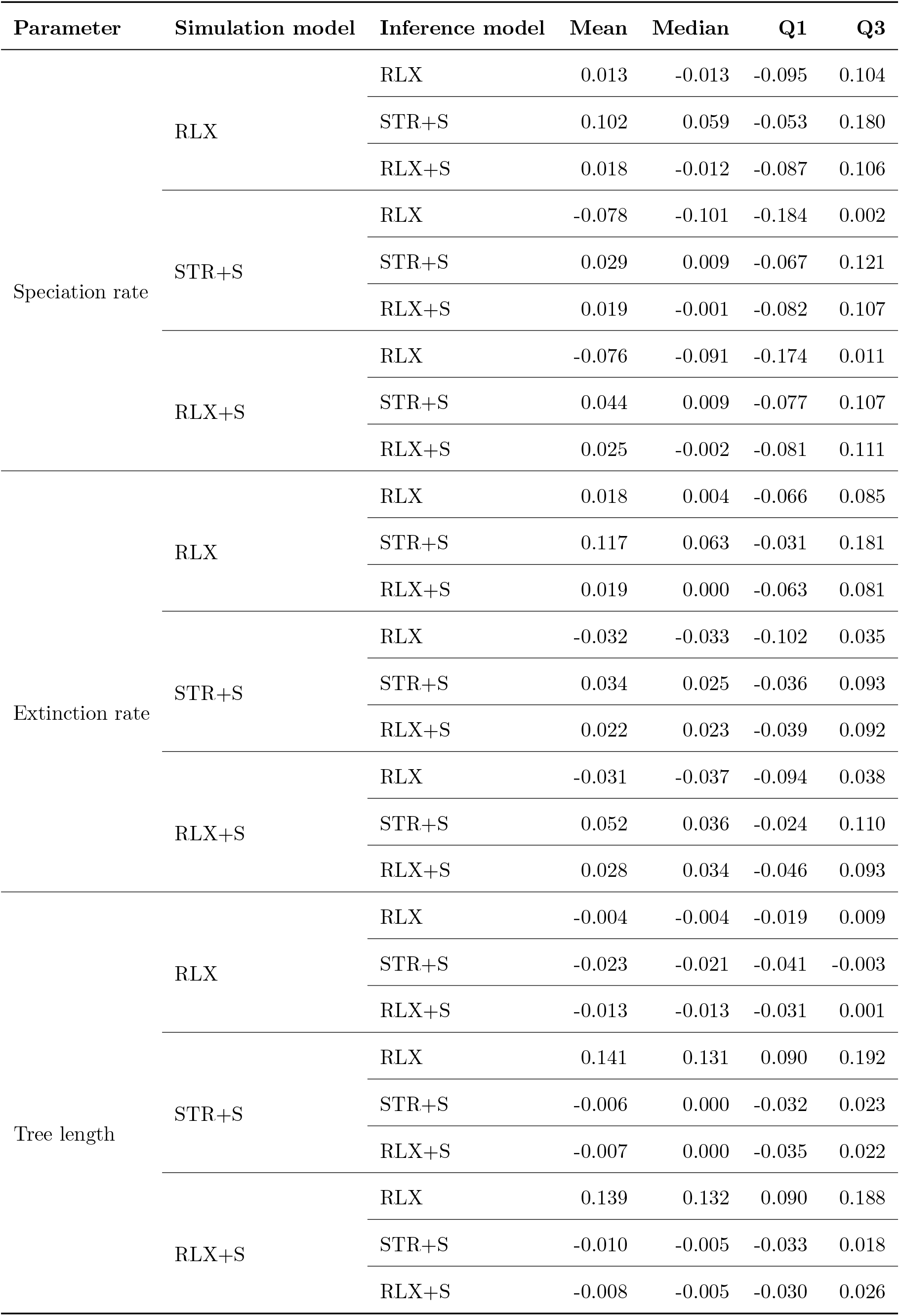
Relative error of posterior median of speciation rate, extinction rate, and tree length. Q1 = 25th percentile, Q3 = 75th percentile. Scenario: complete sequences.

| Parameter | Simulation model | Inference model | Mean | Median | Q1 | Q3 |
| --- | --- | --- | --- | --- | --- | --- |
| Speciation rate | RLX | RLX | 0.013 | -0.013 | -0.095 | 0.104 |
|  |  | STR+S | 0.102 | 0.059 | -0.053 | 0.180 |
|  |  | RLX+S | 0.018 | -0.012 | -0.087 | 0.106 |
|  | STR+S | RLX | -0.078 | -0.101 | -0.184 | 0.002 |
|  |  | STR+S | 0.029 | 0.009 | -0.067 | 0.121 |
|  |  | RLX+S | 0.019 | -0.001 | -0.082 | 0.107 |
|  | RLX+S | RLX | -0.076 | -0.091 | -0.174 | 0.011 |
|  |  | STR+S | 0.044 | 0.009 | -0.077 | 0.107 |
|  |  | RLX+S | 0.025 | -0.002 | -0.081 | 0.111 |
| Extinction rate | RLX | RLX | 0.018 | 0.004 | -0.066 | 0.085 |
|  |  | STR+S | 0.117 | 0.063 | -0.031 | 0.181 |
|  |  | RLX+S | 0.019 | 0.000 | -0.063 | 0.081 |
|  | STR+S | RLX | -0.032 | -0.033 | -0.102 | 0.035 |
|  |  | STR+S | 0.034 | 0.025 | -0.036 | 0.093 |
|  |  | RLX+S | 0.022 | 0.023 | -0.039 | 0.092 |
|  | RLX+S | RLX | -0.031 | -0.037 | -0.094 | 0.038 |
|  |  | STR+S | 0.052 | 0.036 | -0.024 | 0.110 |
|  |  | RLX+S | 0.028 | 0.034 | -0.046 | 0.093 |
| Tree length | RLX | RLX | -0.004 | -0.004 | -0.019 | 0.009 |
|  |  | STR+S | -0.023 | -0.021 | -0.041 | -0.003 |
|  |  | RLX+S | -0.013 | -0.013 | -0.031 | 0.001 |
|  | STR+S | RLX | 0.141 | 0.131 | 0.090 | 0.192 |
|  |  | STR+S | -0.006 | 0.000 | -0.032 | 0.023 |
|  |  | RLX+S | -0.007 | 0.000 | -0.035 | 0.022 |
|  | RLX+S | RLX | 0.139 | 0.132 | 0.090 | 0.188 |
|  |  | STR+S | -0.010 | -0.005 | -0.033 | 0.018 |
|  |  | RLX+S | -0.008 | -0.005 | -0.030 | 0.026 |

**Table S12:** Relative error of posterior median of speciation rate, extinction rate, and tree length. Q1 = 25th percentile, Q3 = 75th percentile. Scenario: 1% fossil sequences.

| Parameter | Simulation model | Inference model | Mean | Median | Q1 | Q3 |
| --- | --- | --- | --- | --- | --- | --- |
| Speciation rate | RLX | RLX | 0.048 | 0.011 | -0.088 | 0.137 |
|  |  | STR+S | 0.144 | 0.096 | -0.032 | 0.257 |
|  |  | RLX+S | 0.065 | 0.025 | -0.087 | 0.161 |
|  | STR+S | RLX | -0.071 | -0.080 | -0.191 | 0.016 |
|  |  | STR+S | 0.046 | 0.036 | -0.111 | 0.159 |
|  |  | RLX+S | 0.020 | 0.012 | -0.107 | 0.105 |
|  | RLX+S | RLX | -0.078 | -0.107 | -0.194 | -0.028 |
|  |  | STR+S | 0.037 | 0.006 | -0.089 | 0.131 |
|  |  | RLX+S | 0.008 | -0.018 | -0.095 | 0.076 |
| Extinction rate | RLX | RLX | 0.052 | 0.017 | -0.083 | 0.132 |
|  |  | STR+S | 0.170 | 0.088 | -0.049 | 0.315 |
|  |  | RLX+S | 0.070 | 0.034 | -0.057 | 0.150 |
|  | STR+S | RLX | -0.036 | -0.043 | -0.132 | 0.039 |
|  |  | STR+S | 0.047 | 0.031 | -0.057 | 0.154 |
|  |  | RLX+S | 0.013 | 0.000 | -0.073 | 0.076 |
|  | RLX+S | RLX | -0.043 | -0.038 | -0.141 | 0.034 |
|  |  | STR+S | 0.040 | 0.038 | -0.048 | 0.121 |
|  |  | RLX+S | 0.002 | 0.000 | -0.077 | 0.089 |
| Tree length | RLX | RLX | -0.008 | -0.008 | -0.067 | 0.048 |
|  |  | STR+S | -0.012 | -0.015 | -0.078 | 0.038 |
|  |  | RLX+S | -0.014 | -0.019 | -0.090 | 0.040 |
|  | STR+S | RLX | 0.128 | 0.128 | 0.046 | 0.205 |
|  |  | STR+S | 0.000 | 0.007 | -0.059 | 0.066 |
|  |  | RLX+S | -0.014 | -0.007 | -0.071 | 0.041 |
|  | RLX+S | RLX | 0.138 | 0.134 | 0.052 | 0.228 |
|  |  | STR+S | 0.006 | 0.012 | -0.063 | 0.067 |
|  |  | RLX+S | -0.003 | 0.002 | -0.064 | 0.062 |

**Table S13:** Relative error of posterior median of speciation rate, extinction rate, and tree length. Q1 = 25th percentile, Q3 = 75th percentile. Scenario: extant samples only.

| Parameter | Simulation model | Inference model | Mean | Median | Q1 | Q3 |
| --- | --- | --- | --- | --- | --- | --- |
| Speciation rate | RLX | RLX | 0.076 | 0.050 | -0.081 | 0.155 |
|  |  | STR+S | 0.207 | 0.140 | -0.010 | 0.321 |
|  |  | RLX+S | 0.104 | 0.078 | -0.065 | 0.211 |
|  | STR+S | RLX | -0.097 | -0.119 | -0.231 | -0.013 |
|  |  | STR+S | 0.046 | 0.018 | -0.101 | 0.155 |
|  |  | RLX+S | 0.034 | 0.013 | -0.111 | 0.119 |
|  | RLX+S | RLX | -0.101 | -0.143 | -0.229 | -0.020 |
|  |  | STR+S | 0.050 | 0.017 | -0.098 | 0.144 |
|  |  | RLX+S | 0.038 | 0.004 | -0.108 | 0.130 |
| Extinction rate | RLX | RLX | 0.103 | 0.056 | -0.073 | 0.214 |
|  |  | STR+S | 0.269 | 0.143 | 0.006 | 0.431 |
|  |  | RLX+S | 0.140 | 0.083 | -0.041 | 0.255 |
|  | STR+S | RLX | -0.103 | -0.139 | -0.220 | 0.022 |
|  |  | STR+S | 0.040 | 0.031 | -0.089 | 0.138 |
|  |  | RLX+S | 0.024 | 0.012 | -0.092 | 0.108 |
|  | RLX+S | RLX | -0.108 | -0.139 | -0.228 | -0.008 |
|  |  | STR+S | 0.043 | 0.026 | -0.085 | 0.146 |
|  |  | RLX+S | 0.027 | 0.019 | -0.090 | 0.126 |
| Tree length | RLX | RLX | -0.013 | -0.012 | -0.046 | 0.024 |
|  |  | STR+S | -0.023 | -0.013 | -0.059 | 0.014 |
|  |  | RLX+S | -0.021 | -0.017 | -0.056 | 0.016 |
|  | STR+S | RLX | 0.109 | 0.085 | 0.007 | 0.215 |
|  |  | STR+S | -0.024 | -0.040 | -0.125 | 0.060 |
|  |  | RLX+S | -0.018 | -0.029 | -0.112 | 0.069 |
|  | RLX+S | RLX | 0.112 | 0.104 | 0.009 | 0.209 |
|  |  | STR+S | -0.023 | -0.024 | -0.119 | 0.046 |
|  |  | RLX+S | -0.020 | -0.029 | -0.109 | 0.042 |

**Table S14:**
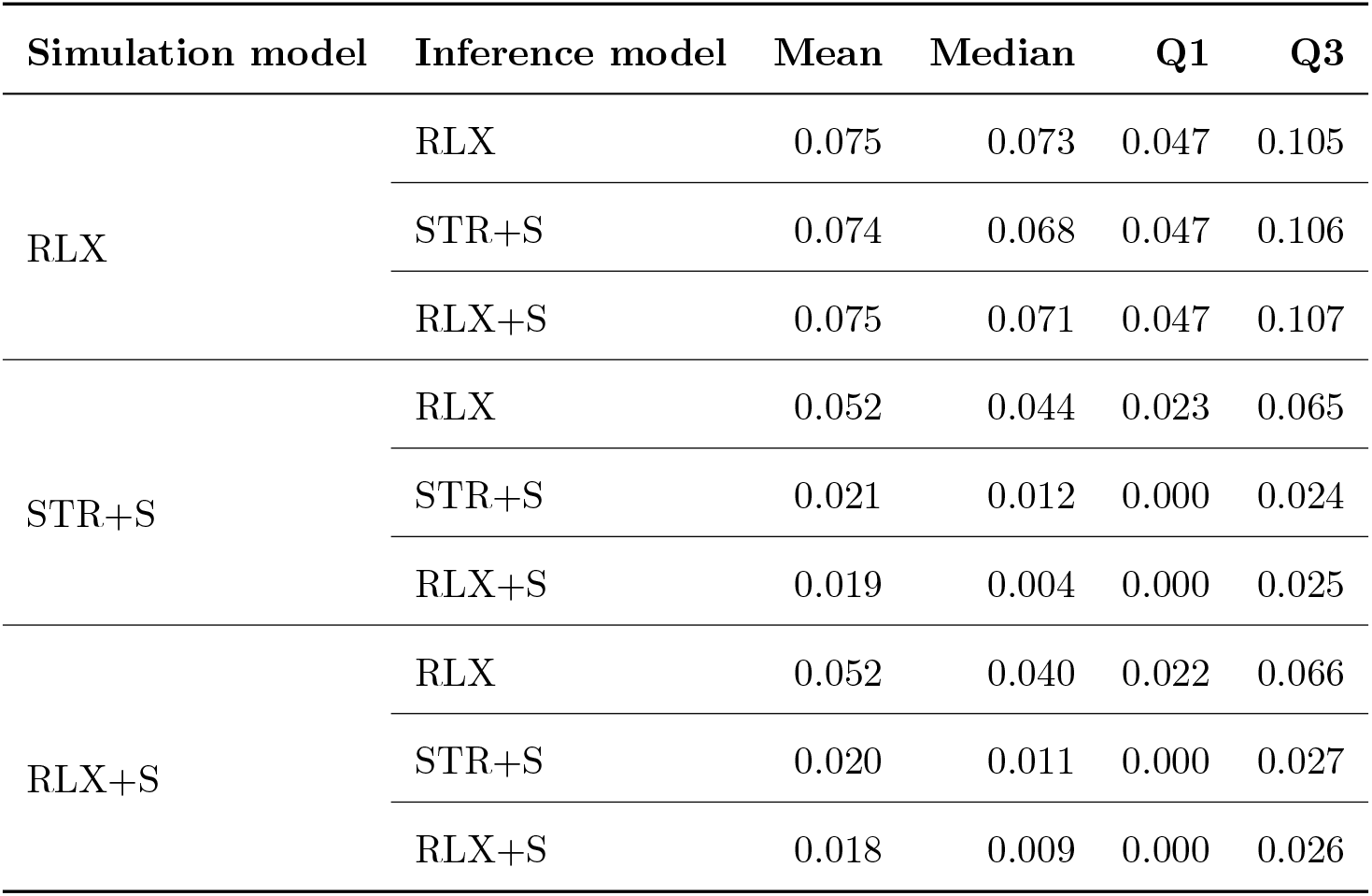
Median normalized Robinson-Foulds distance of posterior trees. Q1 = 25th percentile, Q3 = 75th percentile. Scenario: complete sequences.

| Simulation model | Inference model | Mean | Median | Q1 | Q3 |
| --- | --- | --- | --- | --- | --- |
| RLX | RLX | 0.075 | 0.073 | 0.047 | 0.105 |
|  | STR+S | 0.074 | 0.068 | 0.047 | 0.106 |
|  | RLX+S | 0.075 | 0.071 | 0.047 | 0.107 |
| STR+S | RLX | 0.052 | 0.044 | 0.023 | 0.065 |
|  | STR+S | 0.021 | 0.012 | 0.000 | 0.024 |
|  | RLX+S | 0.019 | 0.004 | 0.000 | 0.025 |
| RLX+S | RLX | 0.052 | 0.040 | 0.022 | 0.066 |
|  | STR+S | 0.020 | 0.011 | 0.000 | 0.027 |
|  | RLX+S | 0.018 | 0.009 | 0.000 | 0.026 |

**Table S15:**
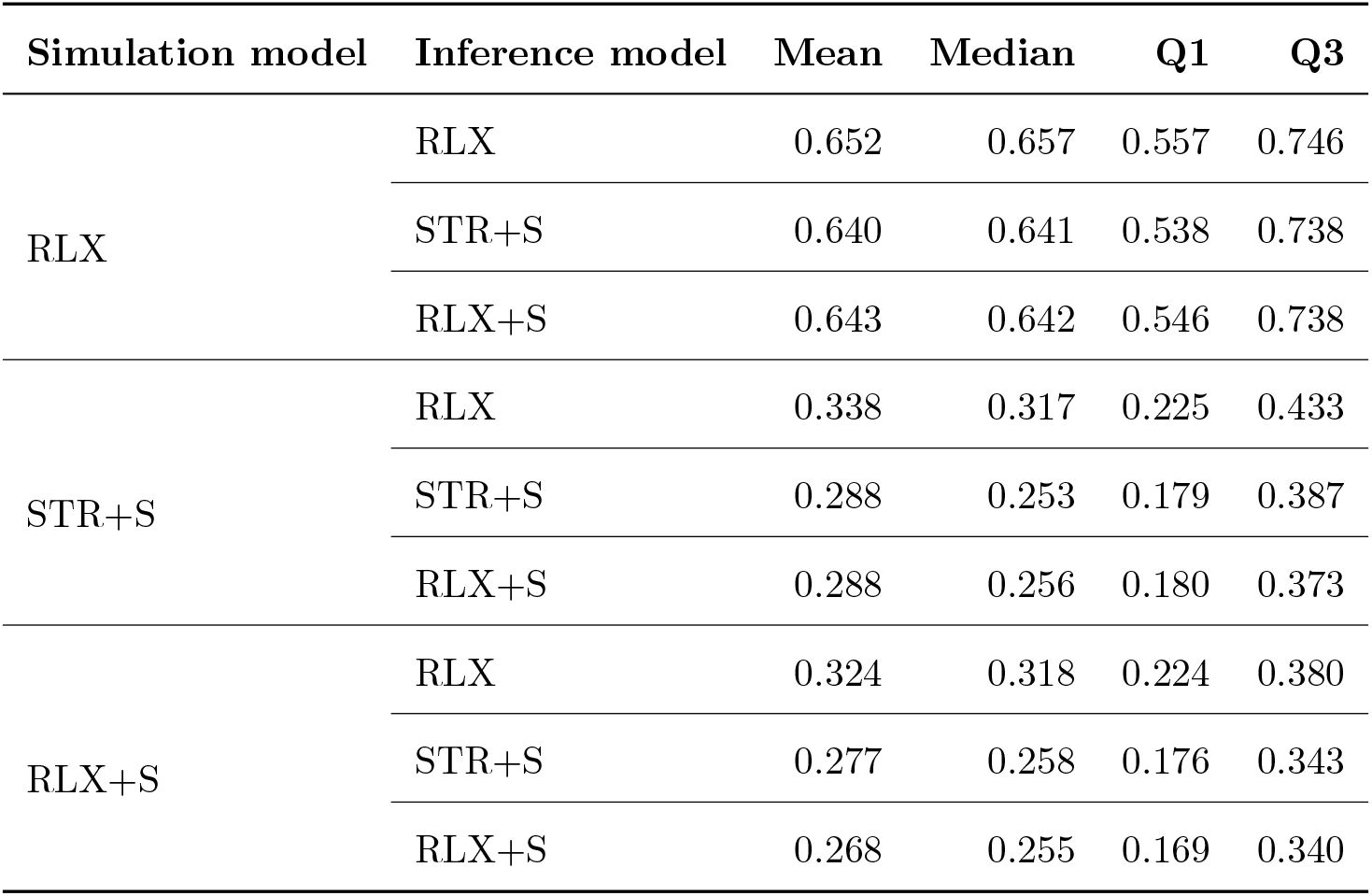
Median normalized Robinson-Foulds distance of posterior trees. Q1 = 25th percentile, Q3 = 75th percentile. Scenario: 1% fossil sequences.

| Simulation model | Inference model | Mean | Median | Q1 | Q3 |
| --- | --- | --- | --- | --- | --- |
| RLX | RLX | 0.652 | 0.657 | 0.557 | 0.746 |
|  | STR+S | 0.640 | 0.641 | 0.538 | 0.738 |
|  | RLX+S | 0.643 | 0.642 | 0.546 | 0.738 |
| STR+S | RLX | 0.338 | 0.317 | 0.225 | 0.433 |
|  | STR+S | 0.288 | 0.253 | 0.179 | 0.387 |
|  | RLX+S | 0.288 | 0.256 | 0.180 | 0.373 |
| RLX+S | RLX | 0.324 | 0.318 | 0.224 | 0.380 |
|  | STR+S | 0.277 | 0.258 | 0.176 | 0.343 |
|  | RLX+S | 0.268 | 0.255 | 0.169 | 0.340 |

**Table S16:** Median normalized Robinson-Foulds distance of posterior trees. Q1 = 25th percentile, Q3 = 75th percentile. Scenario: extant samples only.

| Simulation model | Inference model | Mean | Median | Q1 | Q3 |
| --- | --- | --- | --- | --- | --- |
| RLX | RLX | 0.085 | 0.071 | 0.036 | 0.107 |
|  | STR+S | 0.082 | 0.071 | 0.036 | 0.107 |
|  | RLX+S | 0.083 | 0.071 | 0.036 | 0.107 |
| STR+S | RLX | 0.055 | 0.036 | 0.036 | 0.071 |
|  | STR+S | 0.048 | 0.036 | 0.027 | 0.071 |
|  | RLX+S | 0.047 | 0.036 | 0.000 | 0.071 |
| RLX+S | RLX | 0.055 | 0.036 | 0.036 | 0.071 |
|  | STR+S | 0.050 | 0.036 | 0.000 | 0.071 |
|  | RLX+S | 0.049 | 0.036 | 0.000 | 0.071 |

